# Age-structure as key to delayed logistic proliferation of scratch assays

**DOI:** 10.1101/540526

**Authors:** Ana Victoria Ponce Bobadilla, Thomas Carraro, Helen M. Byrne, Philip K. Maini, Tomás Alarcón

## Abstract

Scratch assays are in-vitro methods for studying cell migration. In these experiments, a scratch is made on a cell monolayer and recolonisation of the scratched region is imaged to quantify cell migration rates. Typically, scratch assays are modelled by reaction diffusion equations depicting cell migration by Fickian diffusion and modelling proliferation by a logistic term. In a recent paper (Jin, W. et al. Bull Math Biol (2017)), the authors observed experimentally that during the early stage of the recolonisation process, there is a disturbance phase where proliferation is not logistic, and this is followed by a growth phase where proliferation appears to be logistic. The authors did not identify the precise mechanism that causes the disturbance phase but showed that ignoring it can lead to incorrect parameter estimates. The aim of this work is to show that a non-linear age-structured population model can account for the two phases of proliferation in scratch assays. The model consists of an age-structured cell cycle model of a cell population, coupled with an ordinary differential equation describing the resource concentration dynamics in the substrate. The model assumes a resource-dependent cell cycle threshold age, above which cells are able to proliferate. By studying the dynamics of the full system in terms of the subpopulations of cells that can proliferate and the ones that can not, we are able to find conditions under which the model captures the two-phase behaviour. Through numerical simulations we are able to show that the resource concentration in the substrate regulates the biphasic dynamics.

## 1 Introduction

Cell migration and proliferation are central processes in the development and homoeostasis of many organisms. They also play a key role in pathological processes such as cancer invasion, chronic inflammatory diseases and vascular diseases [1, 2, 3]. To understand the biochemical and physical cues that regulate cell migration and proliferation, in-vitro assays are used to quantify the ability of cell populations to migrate and proliferate under controlled situations [4, 5, 6]. Scratch assays are typically used to study cell migration [6, 7]. A scratch assay involves: growing a cell monolayer to confluence; creating a “scratch” in the monolayer; and monitoring the cell dynamics as the scratch closes [7]. The resulting time-course data are then analysed to estimate the cell migration rate [8, 9]. Other common cell-based assays are proliferation assays that focus on measuring the cell number or the proportion of cells that are dividing [10]. The simplest proliferation assay consists of growing a cell monolayer to low density on a two-dimensional substrate and measuring the cell number change in time [11, 12, 10]. See Fig. 1 (a) and (b) for a schematic representation of the typical time progression of these two in-vitro assays.

**Fig. 1:**
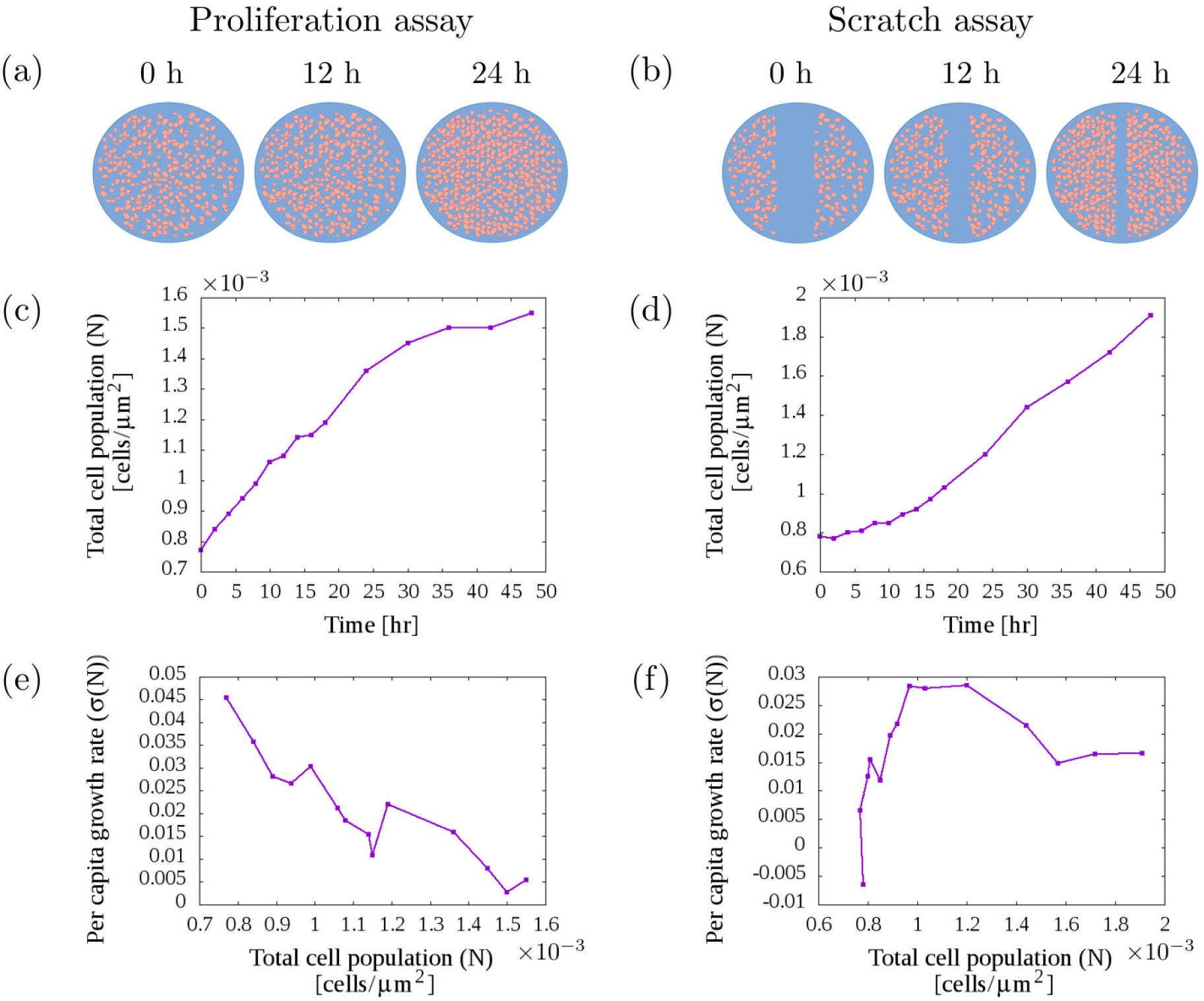
Schematic representation of how the (a) proliferation and (b) scratch assays evolve in time. In the plots (c)-(f), we consider the experimental data from a proliferation and a scratch assay performed in [21]. In (c) and (d) the mean cell population is plotted over the duration of the experiment. In (e) and (f) the per capita growth rate 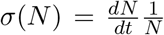 is plotted with respect to the cell population *N*. We calculate the per capita growth rate in the same way as in [21]. A biphasic trend can be observed in (f) (for the scratch assay) but not in (e) (for the proliferation assay).

Many mathematical models have been developed to describe these in-vitro assays and to test hypotheses on the mechanisms that govern cell migration and proliferation [13, 14, 15, 16, 6, 17, 18, 19, 20]. The macroscale dynamics of scratch assays are often described by reaction-diffusion equations [15, 17, 18] in which cell migration is modelled as Fickian diffusion and cell proliferation is modelled by a logistic source term. For the proliferation assay, where no scratch is performed, the logistic growth equation is widely used [14, 16, 6]. Model selection is commonly performed by matching the experimental time-course data with the model evolution and accepting the model if the residual error is small [21, 22, 23]. However, several authors have pointed out that this is not enough for model validation [24, 25].

In [21] the ability of the logistic growth model to describe cell proliferation in scratch assays was studied. A series of scratch and proliferation assays using PC-3 prostate cancer cells were performed and the changes in cell density in two subregions located far from the scratch were quantified. These two subregions were chosen so the changes in cell density were not due to cell migration. To assess the suitability of the logistic growth model, they analysed the model fit to experimental data and the per capita growth rate of the experimental data, 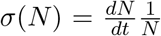, as a function of the cell density, *N*. Calibrating solutions of the logistic model to the experimental data showed a good fit for both assays, however, analysing the per capita growth rate revealed different behaviours between the proliferation and the scratch assays. The authors observed that for proliferation assays, the per capita growth rate could be well described by a linearly decreasing function of the cell density (see Fig. 1 (e)), a result consistent with the logistic model. However, for the scratch assay data, during the first 18 hours of the experiment, the per capita growth rate was found to increase with cell density, whereas for 8 ≤ *t* ≤ 48h, the per capita growth rate was found to decrease approximately linearly with the cell density (see Fig. 1 (f)).

Guided by their experimental observations, the authors in [21] proposed that cell proliferation in scratch assays involves two phases: an initial *disturbance phase* during which proliferation is not logistic and a *growth phase* during which proliferation is approximately logistic. Since this behaviour was not observed in the proliferation assays, the authors concluded that it was caused by the scratching procedure. They hypothesised that scratching might create chemical or mechanical disturbances but did not test their hypothesis.

In this work, we show that a nonlinear age-structured model can account for the disturbance and growth phases observed in scratch assays. We refer to a cell’s *age* as the cell’s temporal position within the cell cycle. The cell cycle consists of four phases: first gap *G*_1_, synthesis stage *S*, second gap *G*_2_ and mitosis stage *M*. *G*_1_ and *G*_2_ consist of growth phases, DNA replication occurs in S phase and cell division in M phase [26]. The progression in the cell cycle is regulated by environmental cues and intracellular checkpoints involving various proteins, in particular cyclins and cyclin dependent kinases [26, 27].

By considering an age-structured model we can investigate how heterogeneity in the cell population age distribution affects the total cell population dynamics. Previous work has shown that heterogeneity in cell age distribution may still generate logistic growth at the population scale [28]. Other age-structured models have described how the cell age distribution can affect the overall cell population dynamics in scratch assays: the speed with which the cells invade the vacant region [29, 30], the efficacy of anti-cancer drugs, particularly phase-specific drugs [31] and the influence of growth factors [13]. Agent-based models have also been considered to study the effect of heterogeneity in cell age distribution on the overall dynamics in in-vitro assays [19, 20].

We consider an age-structured model that was first introduced in [32]. The model considers the interplay between a single cell population and the resource concentration available in the substrate. The model assumes a resource-dependent G1/S transition age, above which, cells are able to proliferate. This critical age naturally divides the cell population into “immature” and “mature” cells. By studying the dynamics of the full system in terms of these subpopulations, we are able to find conditions under which the cell population evolution is of logistic-type and the per capita growth rate follows a biphasic behaviour. We validate our predictions via numerical simulations and show that the resource concentration regulates the disturbance phase.

The remainder of this paper is organised as follows. In Sect. 2, we describe the nonlinear age-structured model. In Sect. 3, we describe the dynamics of the full model in terms of the mature and immature subpopulation dynamics. We then derive conditions under which the per capita growth rate will exhibit biphasic behaviour and logistic-type proliferation. In Sect. 4, we investigate numerically these conditions and determine parameter regimes in which the resource concentration regulates the disturbance phase. We conclude in Sect. 5 with a discussion of the results and conclusions.

## An age-structured model with resource-regulated proliferation

To take into account the cell cycle heterogeneity of a cell population, we consider a McKendrick-von Foerster model [33, 34]. The model considered in this work was first used to describe the mean field dynamics of a stochastic multiscale model of a cell population with oxygen-regulated proliferation [32]. Here the model describes the dynamics of an in-vitro cell population assumed homogeneously distributed in space, like cell cultures in proliferation assays or cells in far away regions from the wound in scratch assays, as in [21]. We are interested in the population dynamics with respect to the resources available in the medium. We denote by *n*(*a, t*), the number of cells of age *a* at time *t*. We consider *a* ∈ [0, ∞]. We denote by *T* > 0 the duration of the experimental observation. The model focuses on the cell population proliferation dependence on the growth factors, oxygen and nutrients available in the medium. We refer to these components as a single, generic *resource* and denote it by *c*(*t*).

We assume cells mature with constant speed 1, die with rate *μ* and proliferate at a rate *b*(*a, c*(*t*)) which we consider to be age and resource-dependent rate [35, 36]. Combining the above assumptions, it is straightforward to show that the evolution of the cell density function *n*(*a, t*) : [0, ∞] × [0, *T*] → ℝ is therefore given by

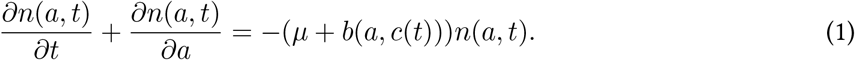

We suppose further that when a cell divides it produces two daughter cells of age *a* = 0. By considering all possible division events, we deduce that

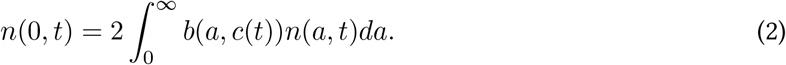

We consider a coarse-grained description of the cell cycle by lumping S, G2 and M into one phase, so we consider a two phase model G1 and S-G2-M. We assume cells proliferate at a constant rate, 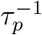, provided they successfully enter the S-G2-M phase. Entry to the S-G2-M phase is regulated by the G1/S checkpoint which has been shown to depend on the presence of resources needed for cell growth [37]. Therefore, we assume that the proliferation rate is given by

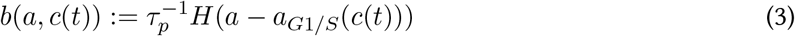

where *H* is the Heaviside function and *a*_*G*1/*S*_ (*c*(*t*)) denotes *the G1/S transition age*, the age at which a cell passes from the G1 to the S phase.

The G1/S transition age, *a*_*G*1/*S*_ (*c*) is given as a function of *c* to capture the dependence of the checkpoint on the available resource concentration. The authors in [32] describe this dependence by a simple scaling which was derived from analysing an oxygen-dependent cell cycle progression model,

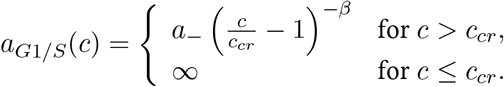

where *a*_−_ and *β* are positive constants. The positive quantity *c*_*cr*_ is the critical resource concentration that allows cell proliferation. For high resource values, the transition age becomes smaller, so the cell requires less time to proliferate. On the other hand, for resource values slightly higher than *c*_*cr*_, the transition age becomes bigger and cells take longer to transition to the G1/S phase and be able to proliferate. For *c* ≤ *c*_*cr*_, cells are assumed to enter the quiescent state.

If we assume that the resource, *c*(*t*), is supplied at a constant rate *S̅* and consumed by all cells at a constant rate *k*; then

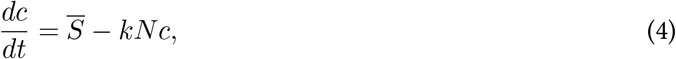

where 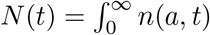 denotes the total number of cells at time *t*.

The full model consists of Eqs. (1) - (4) and the initial conditions

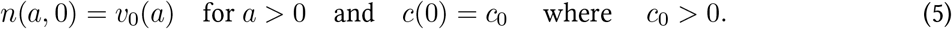

The long-time dynamics of this model was described in [32]. Before beginning our analysis, we summarize below those results from [32] that are relevant for our work.

- For large times, the solution of the age-structured model approaches a separable solution, *Q*(*a, t*) = *A*(*a*) exp(*λt*) i.e. *n*(*a, t*) ≈ *Q*(*a, t*) for *t* ≫ 1, where *λ* satisfies the *Euler-Lotka equation*:

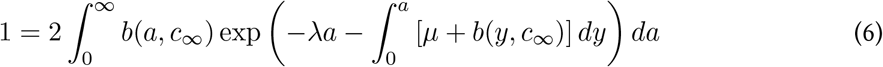

in which *c*_∞_ > 0 is the steady state value of the resource concentration. *A*(*a*) is referred as the *stable age-distribution* of the model. The separable solution is assumed to be in equilibrium when *λ* = 0 and therefore *N* (*t*) → *N*_∞_ and 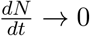 as *t* → ∞.
- For the separable solution to be in equilibrium, the average waiting time to division after the G1/S transition, *τ*_*p*_, must be smaller than the average cell life span, *μ*^−1^.

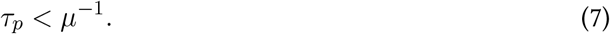
- If Eq. (7) is satisfied, after solving Eq. (6) with *λ* = 0, they obtained that:

– The transition age, *a*_*G*1/*S*_, reaches the steady-state value

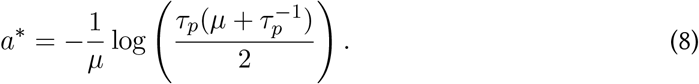
– The steady-state resource concentration is given by

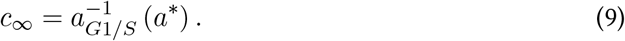
– The steady-state value of the total cell population density is given by:

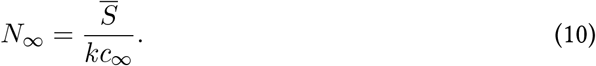

The above results will be useful for parametrising our model.

## 3 Model analysis

In this section, we derive necessary and sufficient conditions for Eqs. (1) - (5) to exhibit logistic-type behaviour and biphasic behaviour in the per capita growth rate.

These conditions can be obtained by distinguishing two subpopulations: cells whose age is greater than the G1/S transition age and which can proliferate, and the cells whose age is below the threshold. We refer to these subpopulations as, respectively, *mature* and *immature* cells. We now analyse the full system with respect to these subpopulation dynamics.

### 3.1 Mature and immature subpopulation dynamics

Let us denote by *X*(*t*) and *Y* (*t*), respectively, the subpopulations of mature and immature cells so that

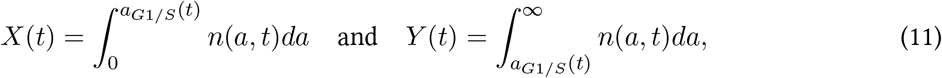

where *a*_*G*1/*S*_ (*t*) := *a*_*G*1/*S*_ (*c*(*t*)).

With these definitions, it is possible to obtain the following result:

#### Theorem 1.

Let X and Y be defined as in (11) and let n(a, t) and c(t) satisfy the model Eqs. (1) - (5). Then the dynamics of X and Y are given by the following system:

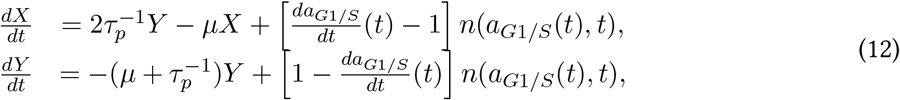

in which 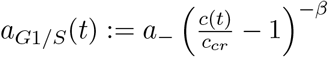 and c(t) is formally given by

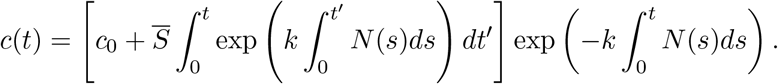

Furthermore, if c(t) ≥ c_cr_ ∀t ∈ [0, T], then the number of cells with G1/S transition age a_G1/S_ (t) at time t, n(a_G1/S_ (t), t), is formally given by

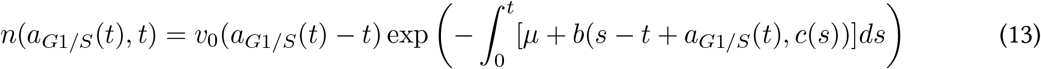

for t ≤ a_G1/S_ (t) and

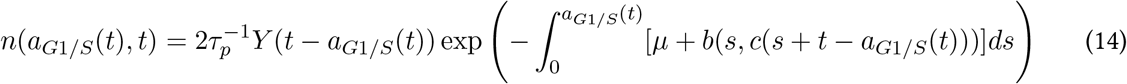

for t > a_G1/S_ (t).

Proof.

Integrating Eq. (1) over the domains [0, *a*_*G*1/*S*_ (*t*)] and [*a*_*G*1/*S*_ (*t*), ∞], we obtain the dynamics of *X* and *Y*, respectively. The evolution of *c*(*t*) is given by solving the differential equation (4) while assuming *N* (*t*) is a known function. Finally, the formula for the cell concentration with G1/S transition age *a*_*G*1/*S*_ (*t*) at time *t*, *n*(*a*_*G*1/*S*_ (*t*), *t*), is obtained by solving the age-structured model by the method of characteristics. The characteristic curves of Eq. (1) are the lines *a* = *t* + *μ* where *μ* is a constant. By solving Eq. (1) along the characteristic curves and taking into account the boundary condition (2) and initial condition (5), we obtain that *n*(*a, t*) is given by:

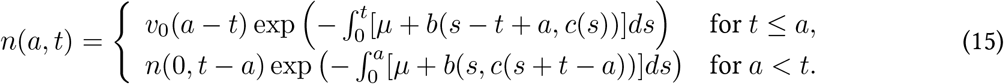

By considering *a* = *a*_*G*1/*S*_ (*t*) and *n*(0, *t* − *a*_*G*1/*S*_ (*t*)) in terms of the mature subpopulation *Y* in Eq. (15), we derive the expressions (13) and (14) for *n*(*a*_*G*1/*S*_ (*t*), *t*).

Theorem 1 reduces the analysis of the full model to the analysis of the system (12). Given Eqs. (13) and (14), we notice that for *t ≤ aG*1/*S* (*t*), (12) consists of a non-homogeneous coupled linear system of differential equations that depends on the initial age-distribution and for *t* > *a*_*G*1/*S*_ (*t*), the dynamics of *Y* is described by a state-dependent delay differential equation,

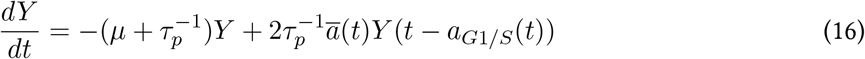

in which 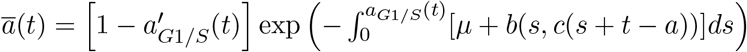.

Having the dynamics of the full model expressed in terms of these subpopulations, we can now describe the overall proliferation and the per capita growth rate in terms of the dynamics of these two subpopulations dynamics. Let us denote by 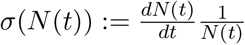, the per capita growth rate. The following theorem gives the evolution of the per capita growth rate and the total population evolution in terms of the mature and immature cell subpopulations.

#### Theorem 2.

The evolution of N (t) and σ(N (t)) for t ∈ [0, T] is given by:

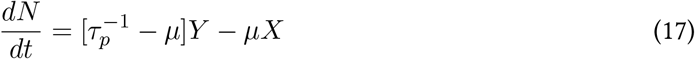

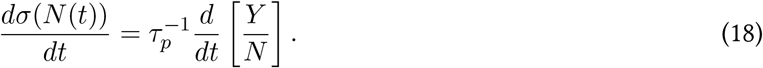

*Proof.* By integrating Eq. (1) in the age domain [0, ∞] we obtain Eq. (17). To derive Eq. (18), we consider the definition of the per capita growth rate and Eq. (17) as follows,

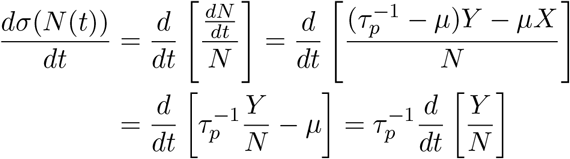

from which we obtain Eq. (18).

### 3.2 Conditions for delayed logistic proliferation

In this section we identify conditions under which the per capita growth rate of the age-structured model exhibits biphasic dynamics and logistic-type growth. First, we interpret the experimental behaviour in mathematical terms:

B1 The cell population undergoes a logistic-type behaviour, i.e,

- The population increases in size monotonically,

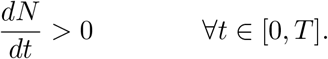
- The population saturates

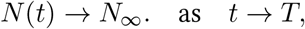

where *N*_∞_ > 0.
B2 The per capita growth rate exhibits biphasic behaviour, i.e. there exists *t*_1_ > 0 such that the per capita growth rate, *σ*(*N* (*t*)), has the following behaviour:

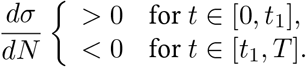

#### Theorem 3.

Necessary and sufficient conditions for the age-structured model with resource-regulated proliferation, Eqs. (1) - (5), to exhibit the behaviour B1 and B2 are as follows

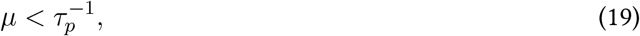

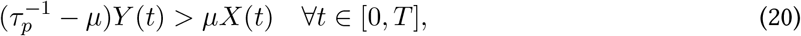

and there exists t_1_ > 0 such that

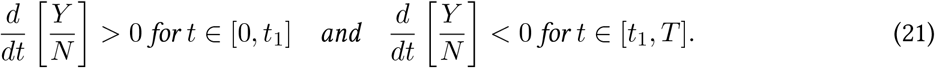

*Proof.* Given the relationship in Eq. (17), it is clear that (20) is true if and only if the derivative of *N* is positive. To derive (21), we notice from the chain rule that 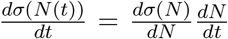 and given that 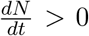, the sign of 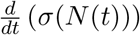 is the same as 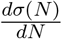. Finally, the condition (19) needs to be fulfilled so that the age-structured model has a stable steady-state solution as described in [32].

Theorem 3 gives necessary and sufficient conditions for the model to exhibit the behaviour described in B1 and B2. We note that the conditions for the biphasic behaviour are given by Eq. (21) which can be interpreted biologically as an initial increase in the fraction of mature cells followed by a stabilization phase in which the mature cell fraction decreases to its steady-state value. Since the fraction of mature cells is regulated by the transition age, *a*_*G*1/*S*_, and this critical age is regulated by the resource concentration, we deduce that changes in the resource dynamics may influence the biphasic behaviour.

## 4 Numerical results

In this section, we perform numerical simulations to verify the predictions of Theorem 3. First, we present the discretization scheme that we use to solve Eqs. (1) - (5).

### 4.1 Discretisation scheme

The model is composed of a hyperbolic partial differential equation with a nonlocal boundary condition coupled to an ordinary differential equation. We use a splitting method to discretise the coupled system. The ordinary differential equation is discretised by an implicit Euler method. The partial differential equation is discretised using the Rothe method: first discretised in time via the implicit Euler method and then in space by linear finite elements [38]. The method is stabilised by considering the streamline upwind/Petrov Galerkin formulation [39]. The finite element discretisation is implemented in C++ using the software DEAL.II [40].

To discretise the age domain [0, ∞], we notice that the cell age is naturally bounded. Hence there exists *a*_*max*_ > 0 such that *n*(*a, t*) = 0 for *a* ∈ (*a*_*max*_, ∞), and so we restrict our attention to *a* ∈ [0, *a*_*max*_]. For the finite element discretisation to be computationally efficient and not create spurious oscillations around the discontinuity point of the proliferation rate (given by the Heaviside function), we discretise the age domain [0, *a*_*max*_] differently in two subdomains: the subdomain [0, *a*^∗^], where *a*^∗^ is given by Eq. (8), is discretised with a step size of *k* = *a*^∗^/2^10^ ≈ 0.00469081, and the domain [*a*^∗^, *a*_*max*_] is discretised with a step size of *k* = (100 *a*^∗^)/2^10^ ≈ 0.0442. We consider a smaller step size for the first domain since for the simulations considered in this work, the evolution of *a*_*G*1/*S*_ (*t*) occurs in this subdomain. For the time discretisation, we consider a time step of *h* = 0.1.

### 4.2 Model parametrisation

We consider as a case study, PC-3 prostate cancer cells. In order to parametrise our model, we use as a guideline the parameters that were estimated in [28] from the experimental time-course data of scratch assays. The authors estimated a proliferation rate of 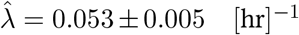. We assume their estimate corresponds in our model to 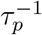, i.e.

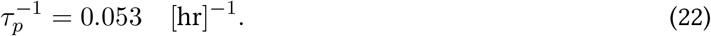

Estimates of doubling times of PC-3 prostate cancer cells are in the range of 25-33 hours [41, 42]. In our model, the doubling time, which we denote as DT, is given by

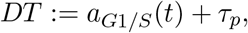

therefore,

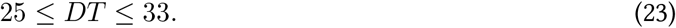

When the cell population reaches the steady-state age distribution, DT is given by:

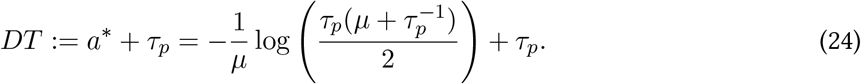

By considering this formula, the estimated value for the proliferation rate 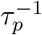 (Eq. (22)) and the range of values for the doubling time (Eq. (23)), we obtain a range of values for the death term, *μ* ∈ [0.0233 − 0.0333]. We assume that *μ* = 0.0283 [hr]^−1^ (the midpoint value).

The carrying capacity was also estimated in [28] they found *K̄* = 2.3 × 10^−3^ ± 2 × 10^−4^ [cells]/[*μm*]^2^. Since the size of the subregion where they estimated *K̄* is 1430 × 200 [*μm*]^2^, the carrying capacity in cell number is

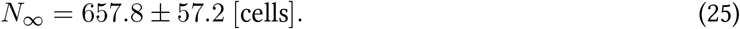

We therefore assume that

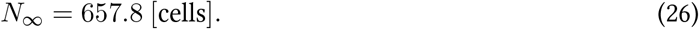

The oxygen consumption rate of PC-3 prostate cancer cells was measured in [43] and found to be approximately 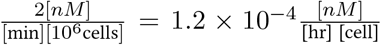. Using this value for the consumption rate, *k*, and the estimate of the carrying capacity (Eq. (26)) in the formula of the steady-state value of the total population in our model (Eq. (10)), we determine the value of the resource flux, 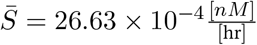.

Regarding the parameters that describe the resource dependence of the G1/S transition age, we consider the parameter values that were derived from an intracellular model for an oxygen-regulated proliferation rate in [32] the dimensionless parameter values are *β* = 0.2 and *c*_*cr*_ = 0.23. For determining the parameter value of *a*_−_, we notice that the admissible range of values for *a*_*G*1/*S*_ (*t*) is 6 ≤ *a*_*G*1/*S*_ (*t*) ≤ 14 given the range values of DT (Eq. (23)) and the estimate of *τ*_*p*_ (Eq. (22)). We consider *a*_−_ = 8.25 [hr], so *a*_*G*1/*S*_ (*t*) is in this range for all our simulations.

We consider the following initial age-distribution that is a multiple of the steady-state age distribution:

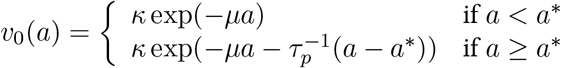

where *a*^∗^ is the steady-state value of the transition age (Eq. (8)) and *κ* is estimated so the initial number of cells corresponds to the average number of cells consider as initial condition in [28] for the most confluent initial condition, *N* (0) = 223. We set *c*_0_ = *c*_∞_ as the steady-state value of the resource concentration given by Eq. (9). The default parameters are listed in Table 1.

**Table 1:** Summary of model parameters

| Parameter | Description | Value | Units | Reference |
| --- | --- | --- | --- | --- |
| $\tau_p^{-1}$ | Proliferation rate | 0.053 | $[\text{hr}]^{-1}$ | [21] |
| $\mu$ | Death rate | 0.028 | $[\text{hr}]^{-1}$ | Estimated |
| $k$ | Consumption rate | $1.2 \times 10^{-4}$ | $[\text{hr}]^{-1}$ | [43] |
| $\bar{S}$ | Resource flux | $26.635 \times 10^{-4}$ | $\frac{[nM]}{[\text{hr}]}$ | Estimated |
| $c_{cr}$ | Critical resource value | 0.023 | [nM] | [32] |
| $\beta$ | Scaling constant | 0.2 | dimensionless | [32] |
| $a_-$ | Scaling constant | 8.25 | [hr] | Assumed |
| $c_0$ | Initial resource value | 0.034 | dimensionless | Estimated |
| $\kappa$ | Scaling constant | 12.488 | dimensionless | Estimated |
| $a_{max}$ | Maximum age | 100 | [hr] | Estimated |

### 4.3 Reference dynamics

Under the default parameter values, the model satisfies the conditions of Theorem 3 and therefore we expect a logistic-type behaviour and a biphasic dynamics. In Fig. 2 (a) and (b) we plot the time evolution of the total cell population *N* (*t*) and of the resource concentration *c*(*t*), respectively. As expected, the total cell population follows a logistic-type behaviour: it grows exponentially and saturates. The resource concentration has an initial increase until it reaches a maximum value and then decreases until it reaches its steady-state value. In Fig. 2 (c) we plot the evolution of the transition age *a*_*G*1/*S*_ and observe its dependence on the resource concentration: it decreases to a minimum value and then increases towards the steady-state value given by Eq. (8). In Fig. 2 (d) we plot the per capita growth rate *σ*(*N*) as a function of the total cell population *N* ; as expected, we observe biphasic behaviour, with an initial increase in *σ* with respect to cell density followed by a linear decrease.

**Fig. 2:**
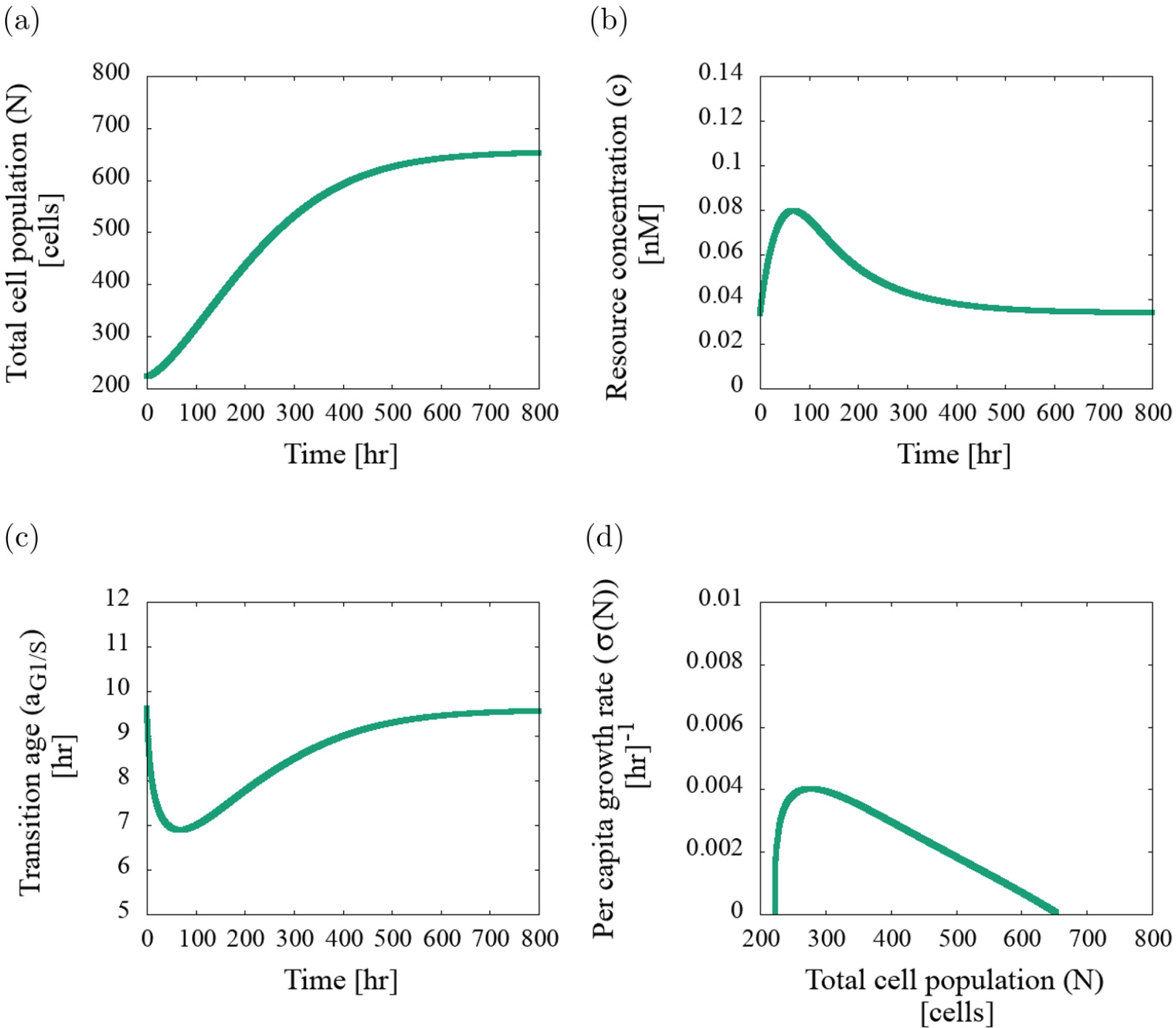
Evolution of the age-structured model with resource-regulated proliferation, Eqs. (1) and (4). In (a) we plot the total cell population evolution and observe it follows a logistic-type growth. In (b) we plot the resource evolution that follows a rapid increase and then a monotonic decrease to the steady-state value. In (c) we plot the evolution of the transition age *a*_*G*1/*S*_ (*t*) and the inverse dependence of the resource concentration on the transition age can be observed. In (d) we plot the per capita growth rate against the total cell population evolution for which two proliferation phases can be observed (a rapid increase, then slower decrease). Parameter values as per Table 1.

### 4.4 Sensitivity analysis

In this subsection we perform a sensitivity analysis focusing on those parameters that modulate the resource dynamics in order to understand their role in determining the biphasic behaviour of the per capita growth rate.

We focus our sensitivity analysis on two parameters that can be experimentally manipulated: the resource flux, *S̅*, and the initial concentration of resource, *c*_0_. We first vary the resource flux, *S̅*, in the domain [25.63 × 10^−4^ − 28.63 × 10^−4^]. We consider this domain so the cell population steady-state value, *N*_∞_, is within the confidence interval given by Eq. (25). As expected, from Eq. (10), the steady-state value at which the total cell population saturates, *N*_∞_, increases as the value of *S̅* increases (Fig. 3). We observe also that the maximum value of *c*(*t*) increases and the minimim values of *a*_*G*1/*S*_ (*t*) decreases as *S̅* increases, see Fig. 3 (c) and (d). We observe in Fig. 3 (d) that the biphasic behaviour is present for all values of the resource flux that were considered.

**Fig. 3:**
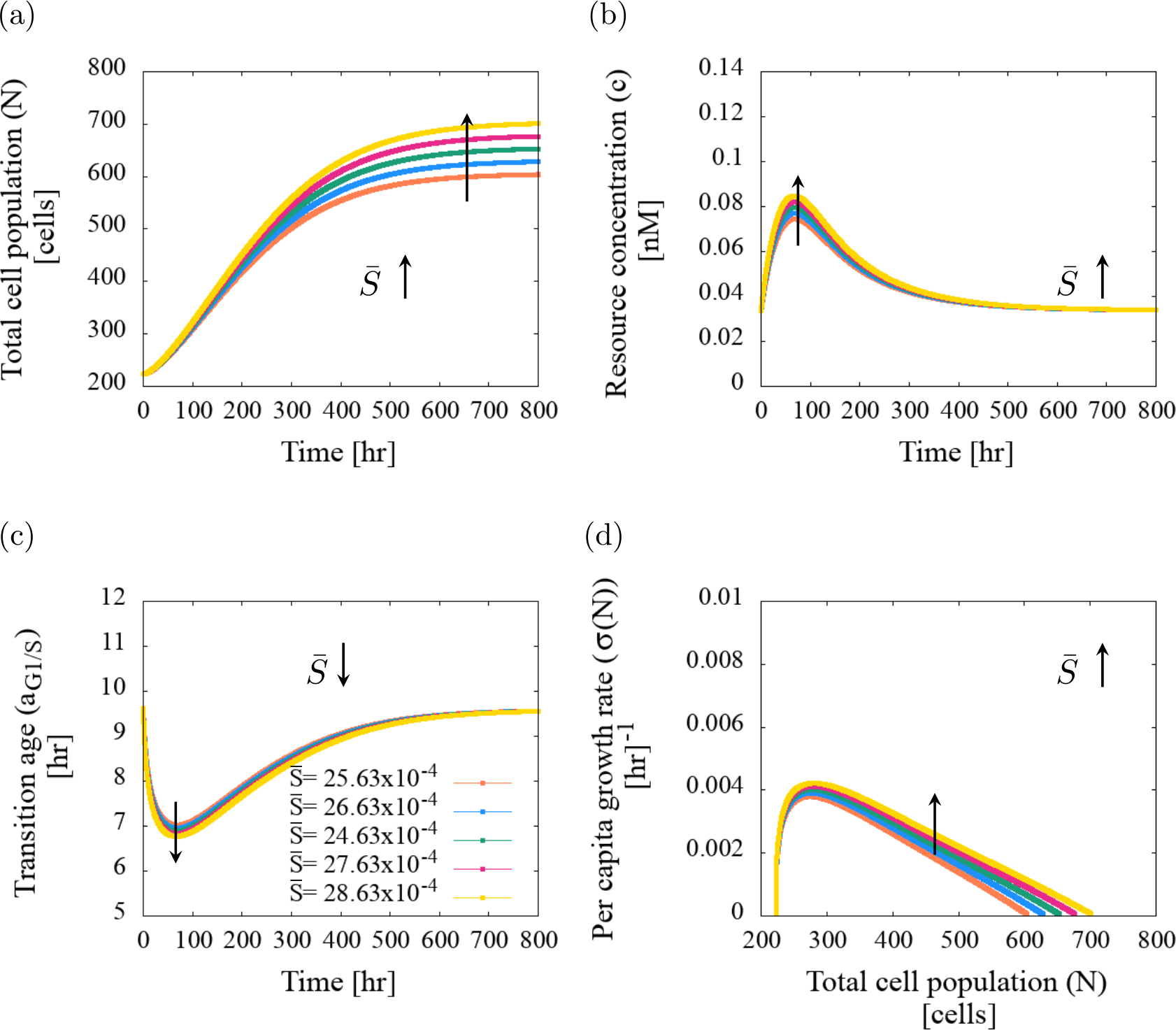
Series of plots showing the dynamics of the coupled age-structured model with resource-regulated proliferation given by Eqs. (1) and (4), as we vary the resource flux, *S̅*. Increasing the value of *S̅*: (a) increases the steady-state value of the total cell population, *N*_∞_, (b) increases the maximum resource concentration and the time it takes to reach the steady-state value, and (c) reduces the minimum value of the transition age *a*_*G*1/*S*_. (d) The plots of the per capita growth rate against the total cell population exhibit biphasic dynamics for the selected values of *S̅*. Parameter values as per Table 1.

We now vary the initial resource concentration *c*_0_. We focus only on increasing the value of the resource concentration above *c*_∞_, since considering values below *c*_∞_ will make the population decrease and this will be in disagreement with experimental observations. In Fig. 4 we show that when we increase *c*_0_, from 0.034 to 0.114, the duration of the disturbance phase is shortened. The model predicts that by increasing *c*_0_, the total cell population evolution does not change significantly, however the resource evolution changes from biphasic to monotonic, i.e. the duration of the disturbance phase decreases and eventually disappears as *c*_0_ increases (see Fig. 4 (b)). Increasing the value of the initial resource concentration shifts the observation time of the dynamics and now it is only possible to observe the resource concentration decline to the steadystate value. This effect can be similarly observed in Fig. 4 (c) where the initial decrease of the transition age *a*_*G*1/*S*_ evolution as *c*_0_ increases. We observe that increasing the initial resource concentration decreases the duration of the disturbance phase where the dependence of the per capita growth rate with respect to the total cell population is not linear (see 4 (d)). We notice that during the disturbance phase, the per capita growth rate does not increase monotonically with respect to the total cell population as we saw previously for the reference dynamics and when varying the resource flux, *S̅*. For the simulations with *c*_0_ bigger than *c*_∞_ = 0.034, the per capita growth rate initially decreases, then increases and finally follows a linear decreasing trend with respect to the total cell population.

**Fig. 4:**
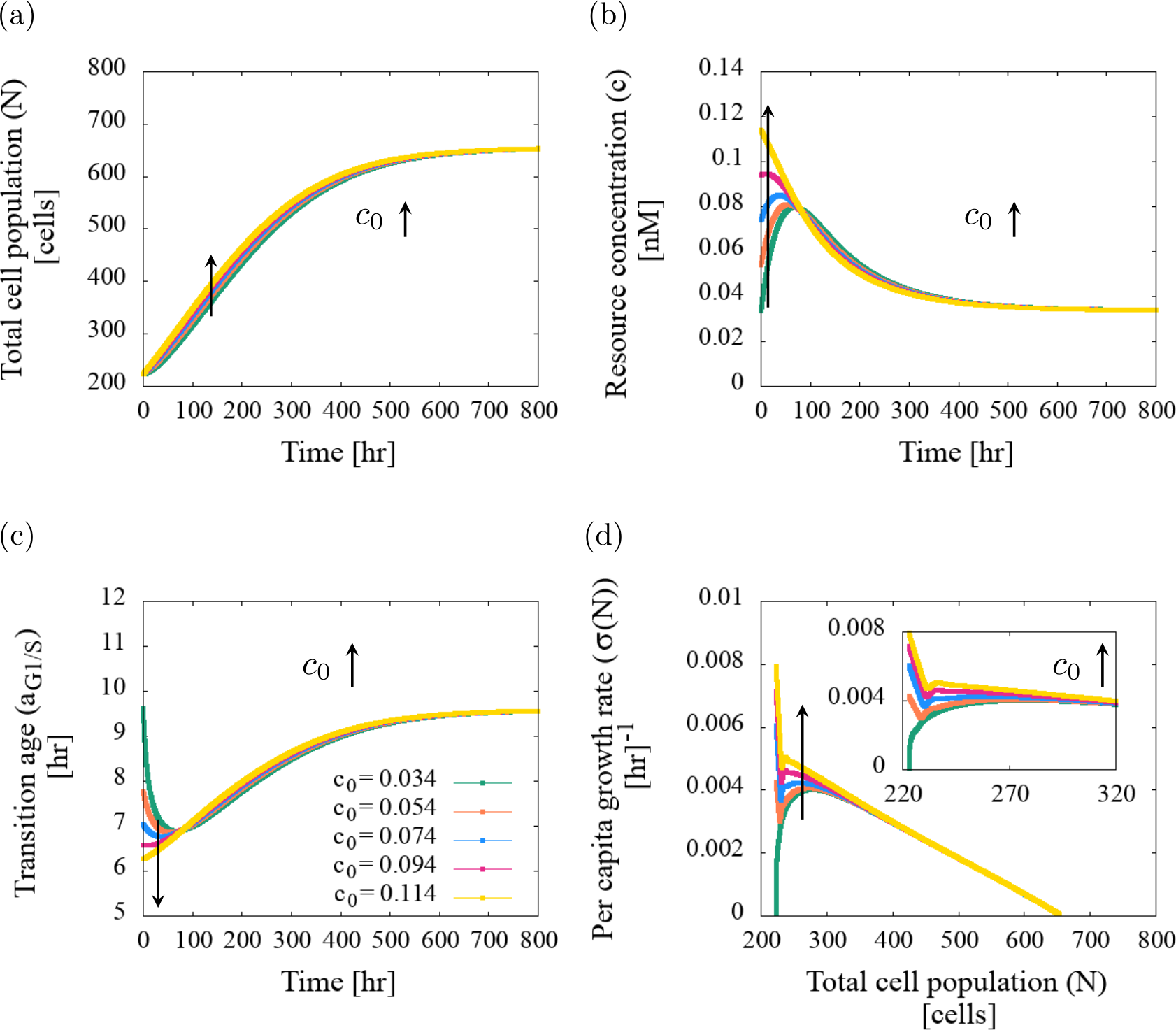
Series of plots showing the dynamics of the coupled age-structured model with resource-regulated proliferation, given by Eqs. (1) and (4), as we vary the initial resource concentration value, *c*_0_. Increasing *c*_0_: (a) does not affect the total cell population, increases the maximum resource concentration but reduces the time it takes to reach it, and (c) makes the initial decrease of the transition age *a*_*G*1/*S*_ disappear. (d) Increasing the initial resource concentration, *c*_0_, affects the plots of the per capita growth rate against the total cell population by shortening the duration of the disturbance phase where there is no monotonic decreasing dependence. Parameter values as per Table 1.

Finally, we perform a sensitivity analysis on the rest of the model parameters and consider different initial age-distributions to show that our model predictions are robust. We observe that uncertainty of the parameters, on the range of values that we considered, has little effect on the overall dynamics. The two proliferation phases are present for all the set of parameters considered (see Supporting Information section 1). We then consider different inital age-distributions that fulfill Theorem 3 and show that the biphasic dynamics is present in all cases considered (see Supporting Information section 2).

## 5 Discussion and conclusions

In this work we have presented an age-structured model with resource-dependent proliferation rate that captures for the first time the biphasic behavior in the per capita growth observed experimentally in [21]. We analysed the full model in terms of two subpopulations: cells which are able to proliferate or not (mature and immature cells). We then derived necessary and sufficient conditions under which the model presents a logistic type behavior and two phases of proliferation: an initial phase, which in [21] was named *disturbance phase,* in which proliferation does not follow a logistic growth and a *growth phase,* where proliferation is approximately logistic. The biphasic behaviour was demonstrated to be a result of an initial increase of the fraction of mature cells followed by a decline to their steady-state concentration. We then parametrised the model using PC-3 prostate cancer cells as a case study. Finally, through numerical simulations, we showed that varying the resource initial value the plot changes the dependence of the per capita growth rate against the total cell population. The model predicts that the duration of the disturbance phase is decreased as the initial resource concentration is increased.

In view of this model, the experimental observations in [21] can be explained as follows: the scratch procedure decreases the cell number in the plate and by replenishing the medium in the same quantity, as customary, the resource concentration is increased, therefore triggering the biphasic dynamics in the per capita growth rate. The predictions of the model can be experimentally tested by modifying the resource concentration in the substrate and examining the plots of per capita growth rate against the total cell population as was performed in [21]. The model predicts that by increasing the initial resource concentration, the disturbance phase would shorten. Varying the resource concentration has been shown to affect the overall dynamics of the scratch assay in other situations [13].

There are different ways that we can extend this work. The model was parametrised using estimates calculated in [21]. Their estimates were calculated with respect to the logistic equation and since we are considering a different model, the values are expected to differ. The model could be parametrised using more appropiate methods as has been done for similar models [44, 13]. Other alternative hypothesis could be analysed: such as mechanical and chemical disturbances [45]. An inference-based modelling approach could be performed to test which model best explains the experimental data or design new experiments to discriminate between feasible alternatives.

In summary, this work identifies the interplay between the mature subpopulation and the resource concentration as the responsibles for the biphasic behaviour observed experimentaly in [21]. Experimental validation and further mathematical modelling will help elucidate the impact of heterogeneity in cell age distribution in the overall dynamics.

## Acknowledgements

AVPB would like to thank Gergely Rost, Maria Vittoria Barbarossa, Wang Jin and Matthew Simpson for helpful discussions. AVPB was supported by the Heidelberg Graduate School of Mathematical and Computational Methods for the Sciences [DFG grant GSC 220 in the German Universities Excellence Initiative].

