## Supporting Information for "Age-structure as key to delayed logistic proliferation of scratch assays"

### S. 1 Parameter sensitivity analysis

We analyse the robustness of the model predictions by performing a parameter sensitivity analysis. We consider as reference, the initial age distribution and the default parameter values given in Section 4.3. Under these conditions, the total cell population follows a logistic-type behaviour and the per capita growth rate exhibits the two proliferation phases, as can be seen in Fig. 2. We analyse if small variations on the parameter values affect the predicted behaviour. When we vary a specific parameter, all other parameters are held fixed at the values reported in Table 1.

We fix the range of values for each parameter by calculating for which values , $550 \leq N_{\infty} \leq 750$, while all the other parameter values are fixed at their values in Table 1. We set the range of values in this way since the carrying capacity estimate of PC-3 prostate cancer cells in [1] is between 600 and 700 cells (see Eq. 25). We extract the following quantities in each simulation:

$$a_{min} := \min_{0 \leq t \leq T} a_{G1/S}(t), \qquad c_{max} := \max_{0 \leq t \leq T} c(t), \qquad \sigma_{max} := \max_{0 \leq t \leq T} \sigma(N(t)),$$

$$t_{amin} \quad \text{such that} \quad a_{G1/S}(t_{amin}) = a_{min}, \quad t_{cmax} \quad \text{such that} \quad c(t_{cmax}) = c_{max},$$

and

$$t_{\sigma max} \quad \text{such that} \quad c(t_{\sigma max}) = \sigma_{max}.$$

In Fig. S1 we plot the quantities that we extract from each simulation when performing the sensitivity analyisis.

After performing the sensitivity analysis, we observe the same qualitative behaviour for all simulations (plots not shown), i.e. the total cell population grows logistically, the resource concentration has an initial increase and then a steady decrease, the transition age has an initial decrease and then a steady decrease, and the two proliferation phases are observed in the plot of the per capita growth rate against the total cell population. In Table S1 we include the mean and variance of the quantities of interest when performing the sensitivity analysis for each of the model parameters. For each parameter, we include the range of values where the parameter was varied, and the mean and the standard deviation of each quantity of interest. We notice that $t_{amin} = t_{cmax}$ so we only include $t_{cmax}$. We observe that in average there is less ten percent variation in the quantities of interest in all the parameter sensitivity analysis, so we can conclude that the parameter uncertainty has little effect on the overall dynamics.

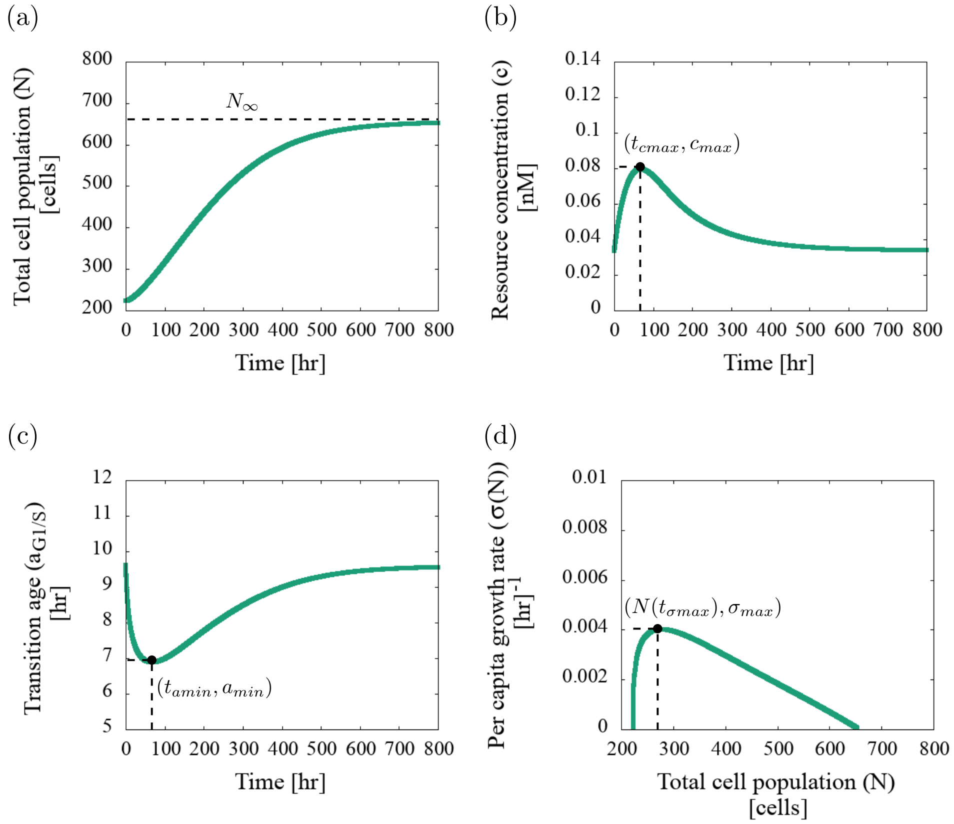


Figure S1: Plot of the quantities extracted from the simulations when performing the sensitivity analysis. The quantities are plotted on top of the model reference dynamics which is introduced on Section 4.3. The quantities of interest are: the resource concentration maximum, $c_{max}$, the transition age minimum, $a_{min}$, the per capita growth rate maximum, $\sigma_{max}$, and the times it takes to reach these values, $t_{cmax}$, $t_{amin}$ and $t_{\sigma max}$, respectively.

Table S1: Results of the parameter sensitivity analysis. In each row, for each model parameter, we specify the range of values in which the parameter was varied and the mean and standard deviation of the quantities of interest.

| Parameter | Range of values | $a_{min}$ | $c_{max}$ | $\sigma_{max}$ | $t_{cmax}$ | $t_{\sigma max}$ |
|---|---|---|---|---|---|---|
| $\tau_p$ | [17.5200,20.2200] | $6.8874 \pm 0.0811$ | $0.0798 \pm 0.0034$ | $4.0510 \times 10^{-3} \pm 1.0061 \times 10^{-3}$ | $68.6200 \pm 8.1282$ | $67.0400 \pm 7.8812$ |
| $\mu$ | [0.0269,0.0291] | $6.8882 \pm 0.1071$ | $0.0799 \pm 0.0044$ | $4.0227 \times 10^{-3} \pm 6.7155 \times 10^{-4}$ | $68.4200 \pm 7.2355$ | $66.1800 \pm 5.7846$ |
| $k$ | [0.0001,0.0001] | $6.8878 \pm 0.1322$ | $0.0800 \pm 0.0055$ | $4.0242 \times 10^{-3} \pm 2.2681 \times 10^{-4}$ | $68.0200 \pm 3.4332$ | $65.8000 \pm 3.3816$ |
| $a_-$ | [7.4829,9.0171] | $6.8847 \pm 0.4419$ | $0.0797 \pm 0.0027$ | $4.0438 \times 10^{-3} \pm 7.6138 \times 10^{-4}$ | $68.2600 \pm 6.1930$ | $65.6200 \pm 5.4961$ |
| $c_{cr}$ | [0.0202,0.0258] | $6.8855 \pm 0.1584$ | $0.0797 \pm 0.0012$ | $4.0287 \times 10^{-3} \pm 2.7157 \times 10^{-4}$ | $67.9200 \pm 2.4035$ | $65.8000 \pm 3.4322$ |
| $\beta$ | [0.1219,0.2781] | $6.9016 \pm 0.3591$ | $0.0797 \pm 0.0011$ | $4.0091 \times 10^{-3} \pm 6.1631 \times 10^{-4}$ | $68.2200 \pm 3.8739$ | $65.5600 \pm 2.2412$ |

### S. 2 Sensitivity analysis of the initial age-distribution

In this section we analyse the impact of varying the initial age-distribution on the model predictions. We consider the initial age-distributions to have the general form:

$$v_0(a) = m \exp(-na), \tag{S1}$$

where $m$ and $n$ are positive real numbers.

In order for the population to initially increase, i.e. $\frac{dN}{dt} > 0$, then $n$ has to satisfy the following inequality,

$$n < \frac{-\log(\mu\tau_p)}{a_{G1/S}(0)}. \tag{S2}$$

Since the initial total cell concentration, $N(0)$, is given by, $N(0) = \frac{m}{n}$, we set the value of $m$ so the initial total cell concentration has the same value as the one we considered in the reference dynamics, $N(0) = 223$ (see Section 4.3). Therefore the value of $m$ is given by

$$m = 223n. \tag{S3}$$

We consider the model parameters as per Table 1. According to Eq. (S2), $n < 0.06642$ We vary $n \in [0.053, 0.065]$ with a step of 0.003 and calculate the value of $m$ through Eq. (S3). For each pair $(m, n)$ we consider the corresponding initial age-distribution and perform a numerical simulation. In Fig. S2 we plot the evolution of different simulations when varying the initial age-distribution. We observe that the total cell population, the resource concentration and the transition age evolution does not change when varying the initial condition. The only plot that changes slightly when varying the initial condition is the plot of the per capita growth rate against the total cell population (see S2 (d)). We observe that for all initial age-conditions considered, the total cell population follows a logistic type behaviour and the plot of the per capita growth rate against the total cell population presents the two proliferation phases.

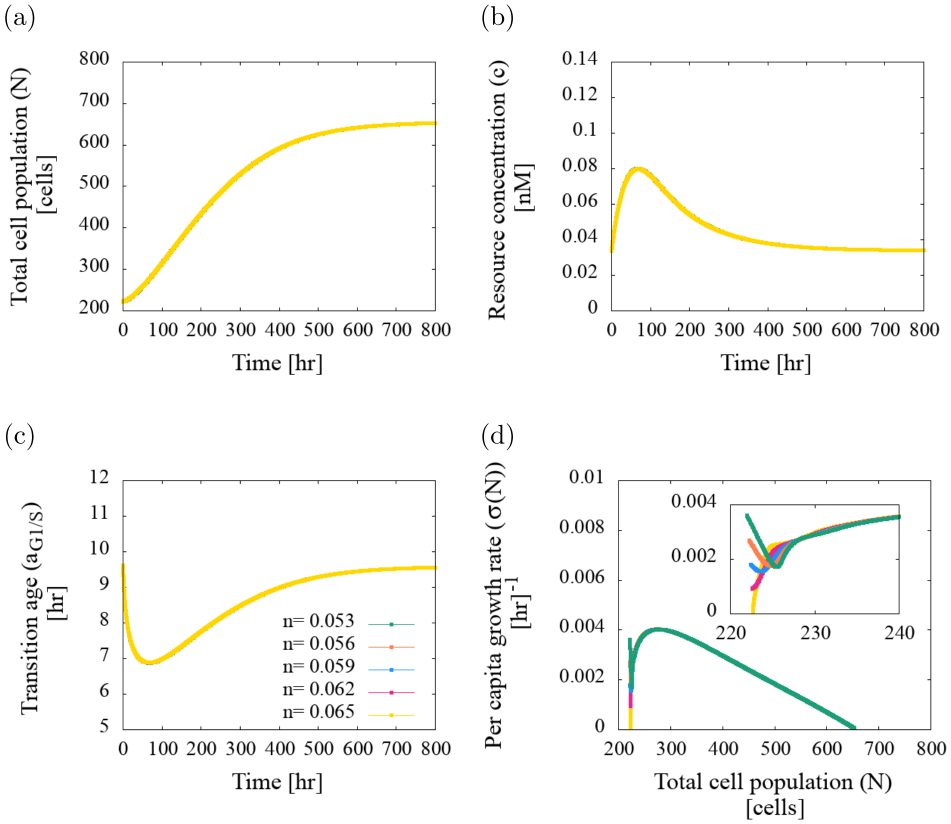


Figure S2: Series of plots showing the dynamics of the coupled age-structured model with resource-regulated proliferation given by Eqs. (1) and (4), as we vary the initial age-distribution. The initial age-distributions have the general form given by Eq. (S1). The value of $m$ is given by Eq. (S3). Varying the initial age-distribution does not affect the: (a) total cell population, (b) resource concentration nor the (c) transtion age evolution. (d) The plots of the per capita growth rate against the total cell population exhibit the two phases of proliferation and the initial dynamics changes slightly as we vary the initial age-distributions. Model parameters are as per Table 1.
